# Temporally Auto-Correlated Predator Attacks Structure Ecological Communities

**DOI:** 10.1101/2021.08.06.455481

**Authors:** Sebastian J. Schreiber

**Affiliations:** Department of Evolution and Ecology and Center for Population Biology, University of California, Davis, CA 95616, USA

## Abstract

For species primarily regulated by a common predator, the *P*\* rule of Holt and Lawton [1993] predicts that the prey species that supports the highest mean predator density (*P*\*) excludes the other prey species. This prediction is re-examined in the presence of temporal fluctuations in the vital rates of the interacting species including predator attack rates. When the fluctuations in predator attack rates are temporally uncorrelated, the *P*\* rule still holds even when the other vital rates are temporally auto-correlated. However, when temporal auto-correlations in attack rates are positive but not too strong, the prey species can coexist due to the emergence of a positive covariance between predator density and prey vulnerability. This coexistence mechanism is similar to the storage effect for species regulated by a common resource. Negative or strongly positive auto-correlations in attack rates generate a negative covariance between predator density and prey vulnerability and a stochastic priority effect can emerge: with non-zero probability either prey species is excluded. These results highlight how temporally auto-correlated species’ interaction rates impact the structure and dynamics of ecological communities.

## Introduction

Predation or resource limitation can regulate populations. When multiple species are regulated by the same limiting factor, long-term coexistence is not expected under equilibrium conditions. Regulation due to a common, limiting resource can result in the *R*\* rule: the species suppressing the resource to the lower equilibrium level excludes other competitors [Volterra, 1928, Tilman, 1990, Wilson et al., 2007]. Regulation due to a common predator can result in the *P*\* rule: the prey species supporting the higher equilibrium predator density excludes the other prey species [Holt and Lawton, 1993, Schreiber, 2004, Holt and Bonsall, 2017]. Yet, many coexisting species share a common resource or a common predator. Understanding mechanisms permitting this coexistence is central to community ecology. One of these coexistence mechanisms, the storage effect, relies on temporal fluctuations in environmental conditions [Chesson and Warner, 1981, Chesson, 1994, Ellner et al., 2016]. Whether an analogous, fluctuation-dependent mechanism exists for species sharing a common predator is studied here.

Similar to species competing for a common resource, species sharing a common predator can exhibit mutually antagonistic interactions Holt [1977]: increasing the density of one prey species leads to an increase in predator density and a resulting increase in predation pressure on the other prey species. Thus, to the uninformed observer, the prey appear to be competing. Empirical support for apparent competition is extensive [Morris et al., 2004, Chaneton and Bonsall, 2000, Holt and Bonsall, 2017] and has significant implications for conservation biology [Gibson, 2006]. When the shared predator is the primary regulating factor, Holt and Lawton [1993] demonstrated that one prey excludes the other via the *P*\* rule. Yet in nature, coexisting species often share common predators. Holt and Lawton [1993] found that spatial refuges, resource limitation, and donor-controlled predation could help mediate coexistence. However, environmentally driven fluctuations in demographic rates did not promote coexistence [Holt and Lawton, 1993, Schreiber, 2021b]. These studies, however, assumed environmental fluctuations are temporally uncorrelated.

In contrast, environmental fluctuations are known, both theoretically and empirically, to mediate coexistence for species competing for a common resource. In a series of influential papers [Chesson and Warner, 1981, Chesson, 1983, 1988, 1994], Chesson identified two fluctuation-dependent coexistence mechanisms: nonlinear averaging and the storage effect. Empirical support for these mechanisms exist in a diversity of systems [Cáceres, 1997, Adler et al., 2006, Angert et al., 2009, Usinowicz et al., 2012, Chu and Adler, 2015, Ellner et al., 2016, Letten et al., 2018]. A key ingredient for the storage effect is a positive covariance between favorable environmental conditions and species’ densities. Temporal auto-correlations, which are commonly observed in environmental factors [Vasseur and Yodzis, 2004, Sun et al., 2018], can generate this positive covariance [Schreiber, 2021a].

Here, temporally auto-correlated fluctuations in demographic rates are shown to mediate coexistence of prey species primarily regulated by a predator and to generate stochastic priority effects. To derive these conclusions, stochastic models of predator-prey interactions are studied using a mixture of analytic and numerical methods.

## Model and Methods

Following Nicholson and Bailey [1935] and Holt and Lawton [1993], the model considers two prey species with densities *N*_1_, *N*_2_ that are regulated by a common predator with density *P*. In the absence of predation, the density of prey *i* increases by a factor, its finite rate of increase *R_i_*(*t*), in generation *t*. Individuals of prey *i* escape predation with probability exp(−*a_i_P*) where *a_i_* is the attack rate on prey i. Captured individuals of prey *i* are converted to *c_i_* predators. To ensure population regulation, predators immigrate at rate *I* > 0 [Holt and Lawton, 1993, Schreiber, 2021b]. Allowing for fluctuations in the demographic rates, the model becomes

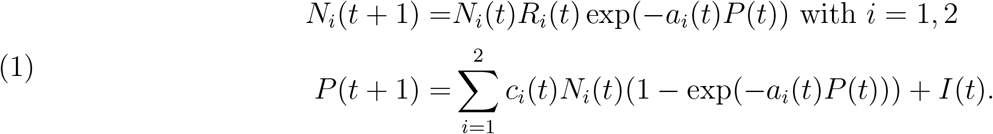

Consistent with meteorological models [Wilks and Wilby, 1999, Semenov, 2008], fluctuations in logarithmic demographic rates are modeled as a first-order auto-regressive processes (see (A1) in Appendix). For example, the log attack rates ln *a_i_*(*t*) are characterized by their means 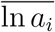, their variances 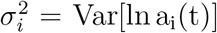, their temporal auto-correlation *ρ* = Cor[ln a_i_(t), ln a_i_(t + 1)], and their cross-correlation *τ* = Cor[ln a_1_(t), ln a_2_(t)]. For the numerical simulations, the auto-regressive processes are Gaussian *i*.e. the attack rates are log-normally distributed.

The dynamics of (1) are explored using analytical and numerical methods. The analytical methods rely on the invasion growth rates (IGRs) of the prey that the correspond to average growth rates when prey species becomes rare [Chesson, 1994, Schreiber, 2000, Schreiber et al., 2011, Grainger et al., 2019, Benaïm and Schreiber, 2019]. When both IGRs for the prey are positive, both species increase when rare and coexist. When both IGRs are negative, there is a stochastic priority effect *i*.e. with non-zero probability either prey species is excluded. When the IGRs have opposite signs, the species with the positive IGR may exclude the other species. Analytical approximations for these IGRs are derived for small environmental fluctuations and computed numerically using R (details in Appendix).

## Results

The invasion growth rates (IGRs) of the prey species are defined by assuming one species, say species *j*, is common (the resident) and the other is infinitesimally rare (the invader), say species *i* ≠ *j*. The the resident species are assumed to coexist (see condition (A2) in Appendix) and have reached a stationary distribution. Let *P_j_*(*t*) be the predator densities at this stationary state. At stationarity, the average intrinsic growth rate of prey *j* equals the average predation rate:

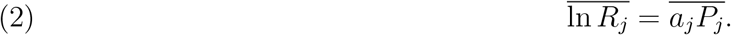

When introduced at (infinitesimally) low densities, the IGR of prey *i* equals the difference between its average intrinsic growth rate and its average predation rate (see (A5) in Appendix)

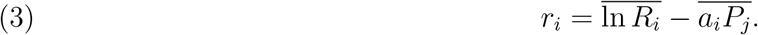

We use equations (2)–(3) to show the *P*\* rule holds when *a_i_*(*t*) are temporally uncorrelated even if the other demographic rates are auto-correlated. Then we show how auto-correlations in *a_i_*(*t*) generate alternative ecological outcomes.

### Temporally uncorrelated attacks and the *P*\*-rule

If the attack rates are temporally uncorrelated (*ρ* = 0), then the average attack rate of the resident prey equals the product of the average attack rate and the average predator density: 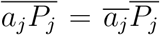. Consequently, equations (2)–(3) imply prey *i*’s IGR is proportional the difference in the average predator densities supported by prey *j* and prey *i*, respectively:

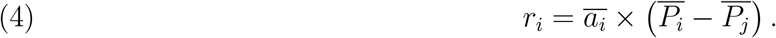

Hence, if prey 1 supports the higher predator density 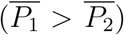, then *r*_1_ > 0 and *r*_2_ < 0 and prey 1 excludes prey 2. The opposite conclusion holds if prey 2 supports the higher predator density 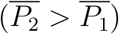.

### The resident and invader attack covariances

Unlike the temporally uncorrelated environments, auto-correlated predator attack rates generate a covariance Cov[a_j_, P_j_] between the predator attack rates *a_j_*(*t*) and the predator densities *P_j_*(*t*) when prey *j* is the resident. This resident attack covariance depends on the auto-correlation *ρ* of *a_j_* (*t*) in a nonlinear fashion (Fig. 1A). An analytical approximation (see equation (A8) in Appendix) shows the resident attack covariance is negative when either the auto-correlation *ρ* is negative (*ρ* < 0) or *ρ* is greater than the reciprocal of the prey’s mean finite rate of increase 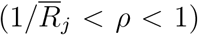. Only for positive, but not too positive auto-correlations 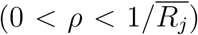 is the resident attack covariance positive.

**Figure 1.**
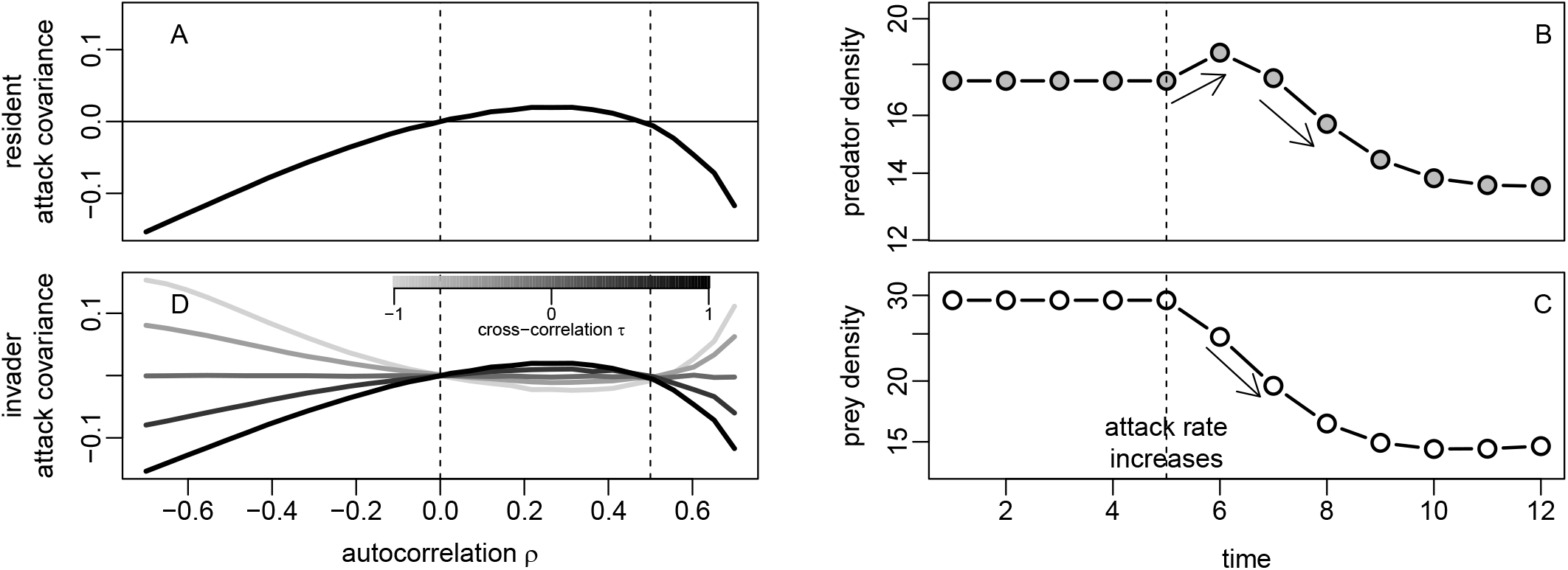
Resident and invader attack covariances depend on auto-correlations in a nonlinear fashion. In (A), the resident attack covariance Cov[a_j_, P_j_] as a function of *ρ* = Cor[ln a_j_(t), ln a_j_(t + 1)]. Dashed lines corresponds to analytical predictions of where this covariance vanishes. This non-linearity stems from the short-term versus long-term effects of an increase in the predator attack rate on the predator (B) and prey (C) densities. In (D), the invader attack covariance Cov[a_i_, P_j_] plotted for different crosscorrelations *τ*. Parameters: 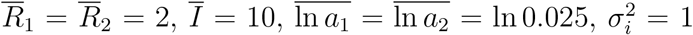, and 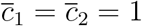.

Figures 1B,C provide a graphical intuition for this non-linear relationship. When the predator’s attack rate increases, there is a short-term increase in the predator’s density. However, the continual decrease in the resident prey’s density ultimately results in a reduction in the predator density. When the auto-correlation is sufficiently positive, increases or decreases in attack rates persist a long time and, thereby, generate a negative resident attack covariance. Less positive auto-correlations or negative auto-correlations play out on shorter time-scales and generate positive or negative covariances, respectively.

Temporally auto-correlated attack rates on prey *i* (the invader) also generate a covariance between their attack rates *a_i_*(*t*) and the predator densities *P_j_*(*t*) (Figure 1D). The sign of this invader attack covariance Cov[a_i_, P_j_] is determined by the cross-correlation *τ* = Corr[ln a_i_, ln a_j_] between the attack rates. When this cross-correlation is positive, the invader and resident attack covariances have the same sign (two darkest lines in Figure 1D). When this cross-correlation is negative, these two covariances have opposite signs (two lightest lines in Figure 1D).

### Auto-correlated attack rates alter ecological outcomes

Due to their effect on the attack co-variances, auto-correlated attack rates can alter ecological outcomes (Figs.2A,B). This impact is best understood when the prey only differ in the timing of predator attacks (i.e. 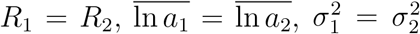 but *τ* = Corr[ln a_i_, ln a_j_] < 1). Then the IGR of prey *i* equals the difference between the resident and invader attack covariances:

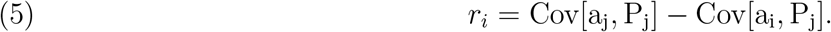

**Figure 2.**
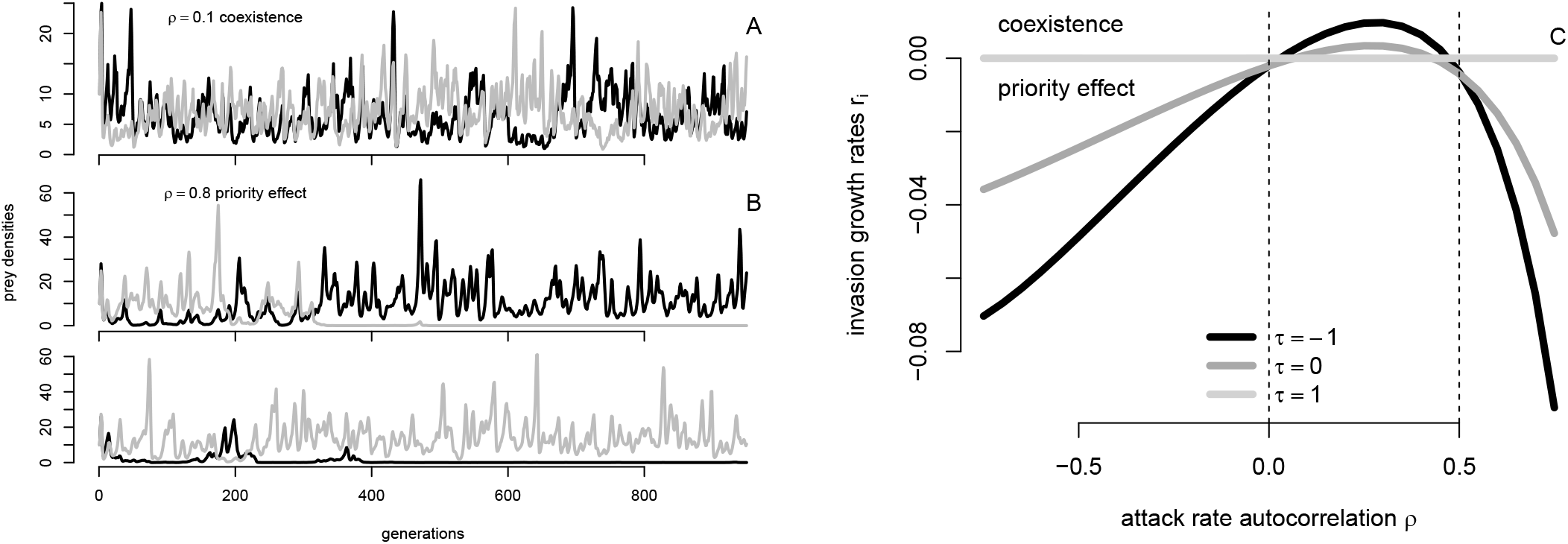
Auto-correlated attack rates alter ecological outcomes. In (A), the dynamics of coexisting prey species and (B) two realizations of the dynamics prey species exhibiting a stochastic priority effect. In (C), the invasion growth rates *r_i_* for both prey species when rare as a function of the temporal auto-correlation *ρ* in attack rates. Different lines correspond to different cross-correlations *τ* = –1, 0,1 in the attack rates. Dashed lines correspond to where the analytic approximation of IGRs *r_i_* are zero. Parameters: 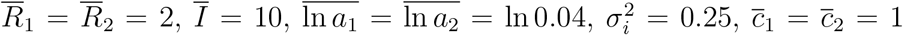, and *τ* = 0.5 for panels (A)–(B).

Whenever the prey species experience differential predation in time (*τ* < 1), the sign of the IGR *r_i_* is determined by the resident attack covariance (Figures 1A,D, 2B). Hence, if the auto-correlation *ρ* in the attack rates is positive but not too positive (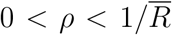, dashed lines in Fig.2C), the IGRs are positive for both species and the prey coexist (Fig.2A). Alternatively, if the auto-correlation ρ is negative or too positive (*ρ* < 0 or 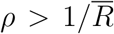), then IGRs are negative for both species and the prey exhibit a stochastic priority effect (Fig.2B).

More generally, when the prey species differ in their intrinsic fitness *R_i_* and differ in their mean attack rates, the invasion growth rate of the prey *i* depends on these differences:

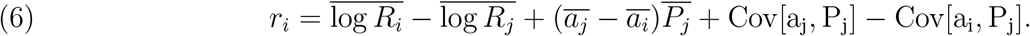

The first term 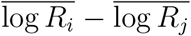 corresponds the difference in the mean intrinsic rates of growth of the two prey species. When prey species *i* has a larger mean intrinsic rate of growth, this term is positive, otherwise it is negative. The second term, 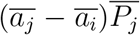, is proportional to the difference in the mean attack rates. When the common prey species is less vulnerable on average (i.e. 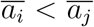), this term is positive, otherwise it is negative. The final pair of terms, the difference in the resident and invader attack covariances, is equivalent to (5). Hence, differences in the prey’s mean intrinsic rates of growth or mean vulnerability to predation can alter the effects of temporally auto-correlated attack rates. For example, large differences in mean attack rates can lead to exclusion despite the attack covariances helping increase IGRs.

## Discussion

For species primarily regulated by a common predator, coexistence isn’t expected under equilibrium conditions: the prey supporting the higher predator density can exclude the other via apparent competition [Holt and Lawton, 1993, Schreiber, 2021b]. Similar to the storage effect for competing species [Chesson, 1994, 2018], we found that environmental fluctuations impacting prey specific attack rates can modify this ecological outcome. Two conditions are necessary for these alternative outcomes. First, fluctuating environmental conditions must differentially impact the predator attack rates on the different prey species. This condition is equivalent to “species-specific responses to environmental conditions” required for the storage effect [Chesson, 1994]. The second condition requires a non-zero, within-generation covariance between predator attack rates and predator density. This attack covariance is analogous to the “environment-competition covariance” of the storage effect for competing species [Chesson, 1994]. When positive, the attack covariance results in relatively lower predation rates on prey that become rare and, thereby, facilitates their recovery from low densities. Hence, coexistence is more likely. When the attack covariance is negative, predation rates are relatively higher on prey that become rare resulting in a stochastic priority effect.

There is empirical evidence that suggests both conditions are likely to occur in nature. For the first condition, differences in prey vulnerability to predation provide multiple pathways for generating asynchronous attack rates among multiple prey [Morin et al., 2021]. These pathways include differences in micro-habitat and refuge availability [Walls, 1995, Woodward and Hildrew, 2002], environmental stressors [Mesa et al., 1994], phenology [Damien and Tougeron, 2019], and morphology and behavior [Riessen et al., 1984, Derting and Cranford, 1989, McPeek, 1990, Einfalt and Wahl, 1997]. For the second condition, temporal auto-correlations are ubiquitous in environmental factors that drive these pathways [Vasseur and Yodzis, 2004, Sun et al., 2018, Di Cecco and Gouhier, 2018] and, as shown here, can generate covariances between attack rates and predator densities.

We demonstrate that both the sign and magnitude of the temporal auto-correlations in attack rates determine the sign of the attack covariance. For positive auto-correlations, the sign of this covariance depends on the time-scale at which the fluctuations occur. When temporal auto-correlations are weak, fluctuations occur on shorter time scales, generate a positive attack covariance, and promote coexistence. In contrast, when temporal auto-correlations are strong, fluctuations occur over longer-time scales, generate a negative attack covariance, and promote stochastic priority effects. Similarly, for continuous-time models of species competing for a common resource, Li and Chesson [2016] found that fast resource depletion generates a positive environment-competition covariance and, thereby, can promote coexistence. Although not stated by Li and Chesson [2016], this positive environment-competition covariance arises from consumer attack rates being positively auto-correlated at the time scale of the resource depletion.

The work presented here and earlier work [Li and Chesson, 2016, Schreiber, 2021a] highlight that co-variances between species densities and per-capita species interaction rates can fundamentally alter the composition and dynamics of ecological communities. These covariances can be driven by the sign and magnitude of auto-correlated fluctuations in environmental conditions. Importantly, the magnitude of these auto-correlations can lead to different ecological outcomes due to differences in transient versus long-term responses of species to changing interaction rates [Bender et al., 1984, Hastings et al., 2018]. Understanding how these effects combine across multiple species, how they interact with other coexistence mechanisms [Chesson, 2018], and how they are impacted by demographic stochasticity [Pande et al., 2020a, Ellner et al., 2020, Pande et al., 2020b] provide significant challenges for future work.

## APPENDIX

This appendix provides the mathematical details for the analysis reported in the main text. With the log demographics rates modeled as first-order auto-regressive processes, the full model is

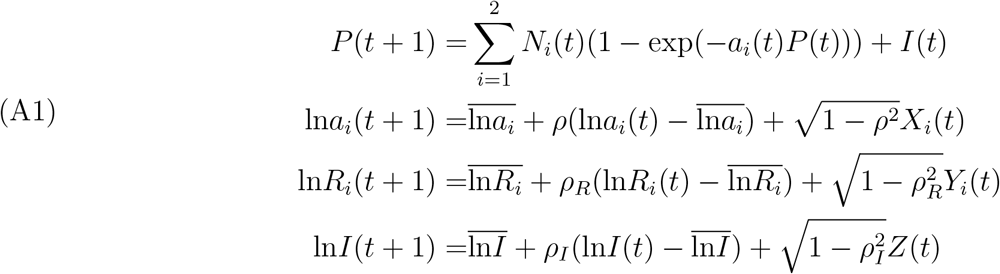

where *X_i_*(1), *X_i_*(2), … are a sequence of i.i.d. random variables with mean 0, variance 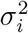 and cross-correlation *τ*, *Y_i_*(1), *Y_i_*(2), … are a sequence of i.i.d. random variables with mean 0 and variance 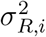 and *Z*(1), *Z*(2), … are a sequence of i.i.d. random variables with mean 0 and variance 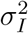. These sequences of i.i.d. random variables are assumed to take values in a compact set. Provided the auto-correlation coefficients are strictly less than one, the processes ln *a_i_*(*t*), ln *R_i_*(*t*), ln *I*(*t*) converge to a compactly supported stationary distribution with means 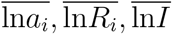 and variances 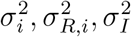.

The dynamics of (A1) are analyzed in four steps. First, using results from Benaïm and Schreiber [2019], I determine under what conditions a single prey species can coexist with the predator. When this condition doesn’t hold, the prey asymptotically goes extinct with probability one. For the remainder of the stochastic analysis, I assume that the condition for each prey species coexisting with the predator holds. Second, I consider the dynamics of the three species model when the predator attack rates are uncorrelated in time; the other demographic rates are allowed to be auto-correlated. Extending the results of Schreiber [2021], I show that the prey species supporting the higher mean predator density excludes the other prey species. Third, I use results from Benaïm and Schreiber [2019] to characterize the general ecological outcomes using the invasion growth rates when rare. Finally, I use a diffusion approximation [Karlin and Taylor, 1981] to derive the approximations for the invasion growth rates *r_i_* presented in the main text.

### Persistence of a single prey species

The proof of the persistence of the single prey species, say species *j*, in the absence of the other prey species closely follows the proof of Theorem 3.1 in [Schreiber, 2021]. The main difference is that [Schreiber, 2021] assumed that the demographic parameters are independent in time i.e. *ρ* = *ρ_I_* = *ρ_R_* = 0. Relaxing this assumption changes the stochastic persistence condition and the expression of the mean predator density. Assume 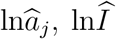, and 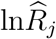 are random variables whose distributions correspond to the stationary distributions of the ln *a_j_*(*t*), ln *R_j_*(*t*), and ln *I*(*t*) auto-regressive processes. The invasion growth rate of prey *j* is given by

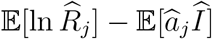

where 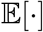 denotes the expectation of a random variable.

Assume that 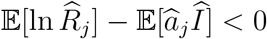. As *P*(*t*) ≥ *I*(*t* – 1) for all *t* ≥ 1, it follows that for all *t* ≥ 1

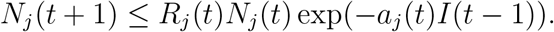

Iterating this inequality implies that

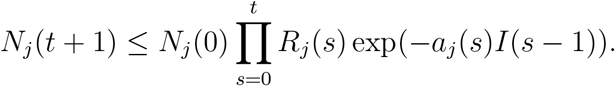

The ergodic theorem for auto-regressive processes implies that with probability one

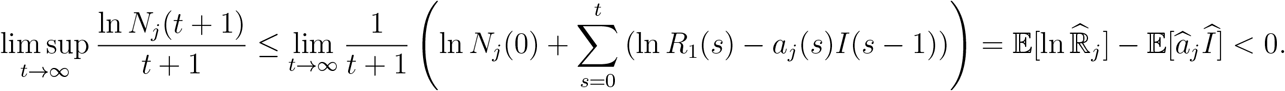

Now assume

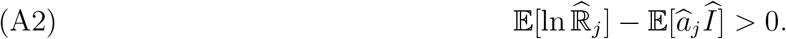

Stochastic persistence of prey *j* follows from [Benaïm and Schreiber, 2019, Theorem 1] where (using the notation for this Theorem) *n* =1, *k* = 4, *X*_1_ = *N_j_*, *Y* = (*P, a_j_*, *I*, *R_j_*), and *p*_1_ = 1.

Still assuming that 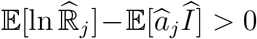, let 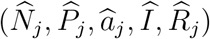 be a random vector that corresponds to a stationary distribution of the model with 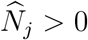 with probability one. Then, [Benaïm and Schreiber, 2019, Proposition 1] implies that the average per-capita growth rate of prey *j* is zero at this stationary distribution i.e.

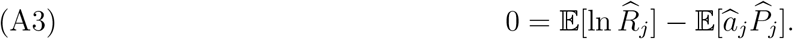

### *P*\* rule when attack rates are serially uncorrelated

Assume that the attack rates are temporally uncorrelated and both prey species can coexist with the predator i.e. 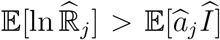 for *j* = 1, 2. Without loss of generality, assume that prey 1 can maintain the higher mean predator density i.e. 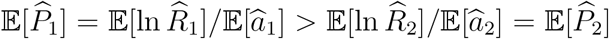. The proof of the exclusion of prey 2 is similar to the proof of Theorem 3.2 in [Schreiber, 2021]. However, [Schreiber, 2021] assumed all the demographic parameters, not only the attack rates, are serially uncorrelated.

First, I show that prey species 1 is stochastically persistent despite the presence of prey species 2. Consider any random vector 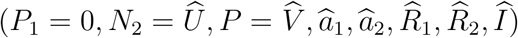 that corresponds to an ergodic, stationary distribution of the model with prey 1 absent. By ergodicity, there are two cases to consider: either 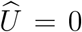 with probability one or 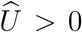 with probability one. In the first case, 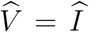 and the invasion growth rate of prey 1 is given by

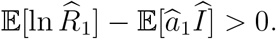

In the second case, stationarity and 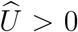 implies the average per-capita growth rate of prey species 2 equals zero i.e.

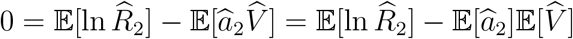

where the second equality follows from *a*_2_(*t*) being i.i.d. in time. Hence,

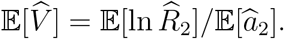

On the other hand, the invasion growth rate of species 1 at this distribution equals

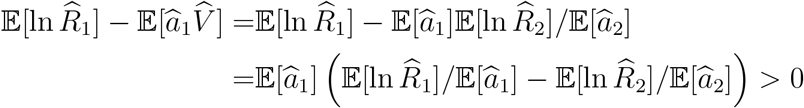

where the last equality follows from the assumption that prey 1 can support the higher mean predator density. Stochastic persistence of prey 1 follows from [Benaïm and Schreiber, 2019, Theorem 1] where (using the notation for this Theorem) *n* =1, *k* = 7, *X*_1_ = *N*_1_, *Y* = (*N*_2_, *P*, *a*_1_, *a*_2_, *I*, *R*_1_, *R*_2_), and *p*_1_ = 1.

Next I show that prey 2 is excluded whenever prey 1 is in the system. Assume *N*_1_(0) > 0 and let *W*(*t*) = (*N*_1_(*t*), *N*_2_(*t*), *P*(*t*), *a*_1_(*t*), *a*_2_(*t*), *I*(*t*), *R*_1_(*t*), *R*_2_(*t*)). Let Π_*t*_(*dw*) be the occupational measure at time *t* i.e. the random measure defined by

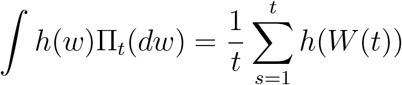

for any continuous bounded function *h*. With probability one, the weak* limit points of Π_*t*_ as *t* → ∞ are an invariant probability measure. Let 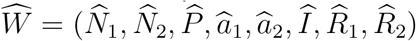 be a random vector whose distribution is given by one of these invariant probability measures. By stochastic persistence of prey 1, 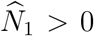 with probability one. Furthermore, by stationarity, the average per-capita growth rate of prey 1 at this distribution equals zero:

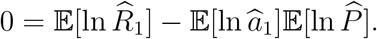

On the other hand, the average per-capita growth rate of prey 2 at this distribution equals

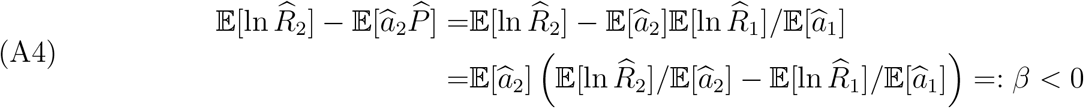

Iterating equation (A1) for *N*_2_ and taking ln implies that

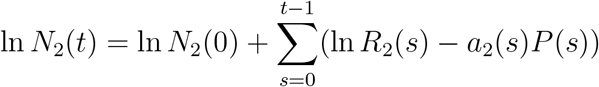

As equation (A4) holds for all weak* limit points of Π_*t*_, it follows that

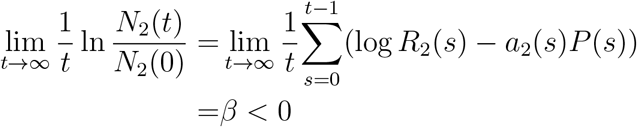

with probability one. Hence, prey species 2 goes extinct asymptotically at an exponential rate.

#### General conditions for coexistence, bistability, and exclusion

When the attack rate is auto-correlated ecological outcomes are characterized the invasion growth rates of both prey species. As before, assume that each prey species can persist with the predator in isolation of the other prey species i.e. 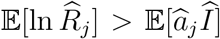 for *j* = 1, 2. Under this assumption, the predator-prey *j* subsystem supports at least one stationary distribution for which *N_j_* > 0 and *N_i_* = 0 with probability one. Let 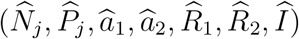 be a random vector whose distribution corresponds to this stationary distribution. Assume this stationary distribution is unique – this happens for example, when the environmental fluctuations are sufficiently large or when the environmental fluctuations are sufficiently small and the predator-prey system without fluctuations has a globally stable equilibrium.

At this stationary distribution, the invasion growth rate of prey *i* is

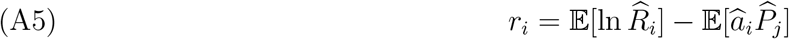

Furthermore, at this stationary distribution, the resident prey *j* growth rate is zero

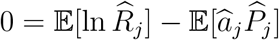

Subtracting this expression from *r_j_* and recalling that 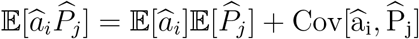 yields

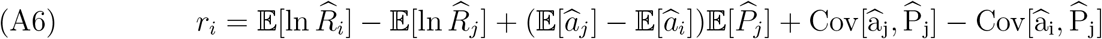

as claimed in the main text.

Benaïm and Schreiber [2019, Theorem 1] implies that if *r*_1_ > 0, *r*_2_ > 0, then the prey coexist in the sense of stochastic persistence. When *r_i_* < 0 and *r_j_* > 0, Benaïm and Schreiber [2019, Corollary 3] implies that species *i* goes exponentially quickly toward extinction when prey *j* is initially present. Finally, when *r*_1_ < 0, *r*_2_ < 0, the same corollary implies that each species is excluded with positive probability whenever the other species is present.

#### Diffusion approximations for prey invasion growth rates

To approximate the invasion growth rates *r_i_* from the previous section, consider the case of small fluctuations in the predator attack rates. Namely, the i.i.d. random variables *X_i_*(*t*) in equation (A1) are given by *X_i_*(*t*) = *εη_i_*(*t*) where *ε* > 0 is small, and *η_i_*(*t*) are i.i.d. with mean zero, variance *v_i_*, and 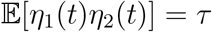 and 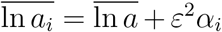 for some choice of *α_i_*. These choices correspond to the classical diffusion scaling [see, e.g., Karlin and Taylor, 1981, Chapter 15]. Furthermore, for simplicity, assume that there are no fluctuations in the intrinsic prey fitness or predator immigration i.e. *Y_i_*(*t*) = *Z*(*t*) = 0 for all time *t* in equation (A1). As in the previous two sections, assume each prey species can persist in the absence of the other i.e. ln 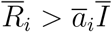 for *i* = 1,2.

In the absence of the fluctuating attack rates, the dynamics of the predator with only prey *j* are given by the deterministic difference equations:

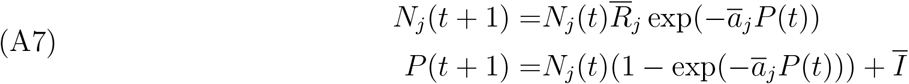

where 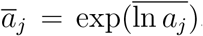. Note that the definition of 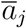 is a slight abuse of notation as 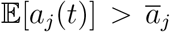 whenever 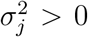 by Jensen’s inequality. For this resident subsystem, the two species coexist about the equilibrium 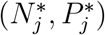 given by

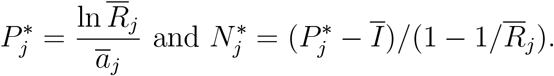

The Jacobian matrix of (A7) at this equilibrium equals

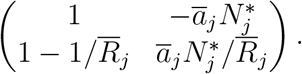

As we are interested in the attack covariance Cov[a_j_(t), P(t)] at stationarity, we also linearize the dynamics of *a*_j_ given by exponentiating the third equation of (A1):

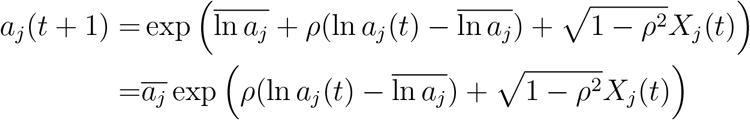

Linearizing with respect to small fluctuations in *a_j_*(*t*) and *X_j_*(*t*) gives the approximation

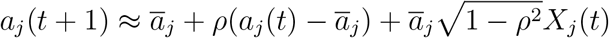

i.e. an autoregressive process where the noise term has an additional weighting term 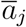. Hence, for small fluctuations (i.e. small *ε* > 0), one can approximate the fluctuations

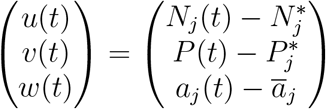

around the equilibrium values as a first-order auto-regressive process:

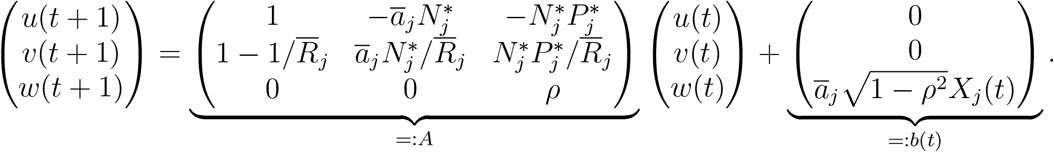

The covariance matrix *C* of 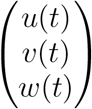 at stationarity [see, e.g., Schreiber and Moore, 2018] satisfies

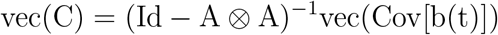

where vec(*) is a column vector given by concatenating the columns of its argument * and ⊗ denotes the Kronecker product. Carrying out this calculation yields the approximation

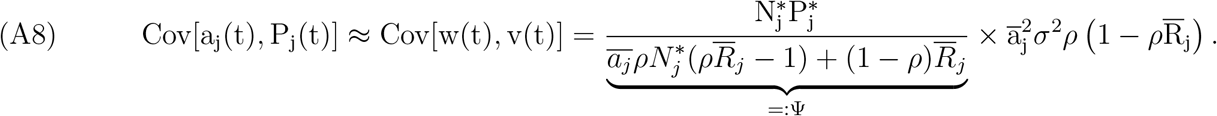

Next I show that Ψ > 0 whenever 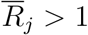. The definition of 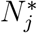 implies that

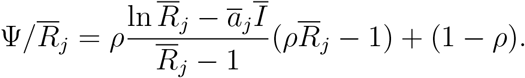

Note that 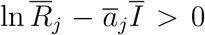 by assumption. For *ρ* ≤ 0 and 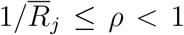, the first term of 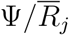 is non-negative and the second term is positive. Hence, Ψ > 0 for *ρ* ≤ 0 and 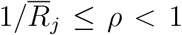. Assume 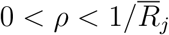. Then

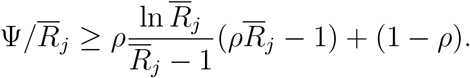

as 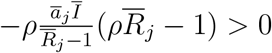. Define *f*(*x*) = ln *x*/(*x* – 1). I claim that *f*(*x*) < 1 for *x* > 1 in which case

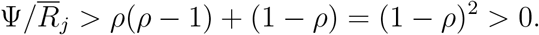

To prove the claim about *f*, l’Hopital’s rule implies that

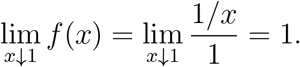

Furthermore,

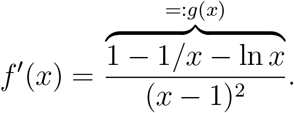

For *x* > 1, the sign of *f*′(*x*) is determined by the sign of *g*(*x*). As *g*(1) = 0 and *g*′(*x*) = (1/*x*)(1/*x* – 1) < 0 for *x* > 1, *f*′(*x*) < 0 for *x* > 1. Hence, as claimed, *f*(*x*) < 1 for *x* > 1.

Equation (A8) and Ψ > 0 implies that the sign of the attack covariance Cov[a_j_(t), P_j_(t)] equals the sign of 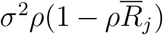. In particular, if 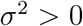, then the attack covariance is positive if 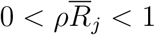 and negative if either *ρ* < 0 or 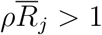 as claimed in the text.

